# A transcription factor module mediating C_2_ photosynthesis

**DOI:** 10.1101/2023.09.05.556297

**Authors:** Patrick J. Dickinson, Sebastian Triesch, Urte Schlüter, Andreas P.M. Weber, Julian M. Hibberd

**Author notes:** Email addresses PD, JMH.

## Abstract

C_4_ photosynthesis has arisen from the ancestral C_3_ state in over sixty lineages of angiosperms. It is widely accepted that an early step in C_4_ evolution is restriction of glycine decarboxylase activity to bundle sheath cells to generate the so-called C_2_ pathway. In C_2_ *Moricandia* species, changes to the *cis*-regulatory region controlling expression of the P-subunit of *GLYCINE DECARBOXYLASE* (*GLDP*) in mesophyll cells enables this trait, but the mechanism underpinning *GLDP* expression in the bundle sheath is not known. We identify a MYC-MYB transcription factor module previously associated with the control of glucosinolate biosynthesis as the basis of *GLDP* expression in bundle sheath cells. In C_3_ *Arabidopsis thaliana* this module drives *GLDP* expression in bundle sheath cells along with as yet unidentified factors driving expression in mesophyll cells. In the C_2_ species *Moricandia arvensis, GLDP* expression is lost from mesophyll cells and the MYC-MYB dependent expression in the bundle sheath is revealed. Evolution of C_2_ photosynthesis is thus associated with a MYC-MYB based transcriptional network already present in the C_3_ state. This work identifies a molecular genetic mechanism underlying the bundle sheath accumulation of glycine decarboxylase required for C_2_ photosynthesis and thus a foundational step in the evolution of C_4_ photosynthesis.

## Introduction

Fixation of CO_2_ during photosynthesis is central to life. In plants this is dependent on Ribulose 1,5-Bisphosphate Carboxylase Oxygenase (RuBisCO) operating as part of the Calvin-Benson-Bassham cycle. However, in addition to reacting with CO_2_ RuBisCO catalyses a side-reaction with O_2_ to produce the toxic metabolite phosphoglycolate (Bowes et al., 1971). The photorespiratory pathway metabolises phosphoglycolate, but CO_2_ is lost and ATP, NADPH and amino acids are required (Tolbert, 1971). As temperatures increase the ratio of oxygenation to carboxylation reactions at the RuBisCO active site increases and so losses from photorespiration become more significant (Jordan and Ogren, 1984). It is widely thought that carbon concentrating mechanisms such as C_4_ photosynthesis evolved to reduce the metabolic costs of photorespiration. In the case of the C_4_ pathway this involves modifications to leaf anatomy, cell biology and biochemistry (Hatch, 1987). Typically, C_4_ biochemistry enables initial fixation of bicarbonate by the enzyme phospho*enol*pyruvate carboxylase in mesophyll cells. Subsequent decarboxylation of C_4_ acids releases high concentrations of CO_2_ in a compartment such as the bundle sheath (Sage, 2001; Christin et al., 2013) and so the oxygenase activity of RuBisCO is reduced (Leegood, 2002; Carmo-Silva et al., 2015).

Some genera contain species that possess biochemical and anatomical characteristics associated with both C_3_ and C_4_ photosynthesis. Such plants have become known as C_3_-C_4_ intermediates or more recently C_2_ species (Sage et al., 2012; Lundgren, 2020), and although statistical modelling predicted that the order of C_4_ trait acquisition is flexible (Williams et al., 2013), a consistent and early event is considered a shift of glycine decarboxylase from mesophyll cells such that its activity is restricted to the bundle sheath (Rawsthorne et al., 1988; Morgan et al., 1993). Repositioning of glycine decarboxylase to the bundle sheath is conjectured to initiate greater rates of CO_2_ release and thus increased photosynthetic activation of this tissue. The two-carbon glycine molecule thus provides CO_2_ for photosynthesis and led to the term C_2_ photosynthesis. Glycine decarboxylase is made up of four subunits and loss of expression of the P-subunit (*GLDP*) from the mesophyll has repeatedly driven the appearance of C_2_ photosynthesis (Rawsthorne et al., 1988; Morgan et al., 1993; Schulze et al., 2016). One example of this is found in the Brassicaceae family where the *Moricandia* genus contains both C_3_ and C_2_ species (Schlüter et al., 2017; Schlüter et al., 2023).

In Brassicaceae species a DNA region referred to as the mesophyll (M) box is highly conserved in promoters of the *GLDP1* gene from C_3_ and C_2_ species (Adwy et al., 2015). Promoter deletion analysis showed that this region is involved in driving expression in mesophyll cells in *A. thaliana* and C_3_ *M. moricandioides* (Adwy et al., 2015, Adwy et al., 2019). Insertion of transposable elements between the M box and the core promoter is thought to abolish mesophyll expression of *GLDP1* in C_2_ species leading to bundle sheath preferential expression (Triesch et al., 2022).

In contrast to our understanding of how loss of mesophyll expression of *GLDP1* is brought about, the molecular architecture enabling the emergence of bundle sheath *GLDP1* expression in the Brassicaceae has not yet been defined. Using C_3_ *Arabidopsis thaliana* we first show that a bipartite MYC and MYB transcription factor module responsible for directing the transcription factor *MYB76* and thus glucosinolate biosynthesis genes to the bundle sheath is also able to pattern *GLDP1* to this tissue. In the C_3_ state this MYC-MYB module operates in parallel with the M box (Adwy et al 2015) to ensure expression in both mesophyll and bundle sheath cells. The MYC-MYB binding sites are conserved in C_2_ *M. arvensis* whereas the insertion of transposable elements has shifted the M box so that it’s function is disrupted (Triesch et al., 2022). Therefore, this MYC-MYB module allows expression of *GLDP1* and assembly of the glycine decarboxylase holoprotein specifically in bundle sheath cells. We thus identify a molecular architecture in the C_3_ state operating in both *cis* and *trans* that underpins a foundational trait associated with the evolution of C_2_ and C_4_ photosynthesis.

## Results and Discussion

### In C_3_ *A. thaliana* MYC and MYB transcription factors drive expression in the bundle sheath which combined with a mesophyll module generates broad expression across the leaf

*A. thaliana* contains two copies of *GLDP*, both of which are expressed in leaves (Supplemental Fig. 1A) (Aubry et al., 2013). All Brassicaceae lineages containing C_2_ species are members of the monophyletic Brassiceae tribe. In the Brassiceae, *GLDP2* is lost leaving *GLDP1* as the only *GLDP* copy, preconditioning the evolution of C_2_ photosynthesis (Schlüter et al., 2017). As *GLDP1* is the copy of *GLDP* involved in C_2_ photosynthesis we focussed on understanding the expression of *GLDP1*. As expected, the *GLDP1* promoter from C_3_ *A. thaliana* drove constitutive expression in leaves (Fig. 1A, Supplemental Fig. 2). Consistent with previous analysis (Adwy et al., 2015), a 5’ deletion removing the M box revealed that the proximal promoter is sufficient to generate expression in bundle sheath strands (Fig. 1B, Supplemental Fig. 3).

**Figure 1.**
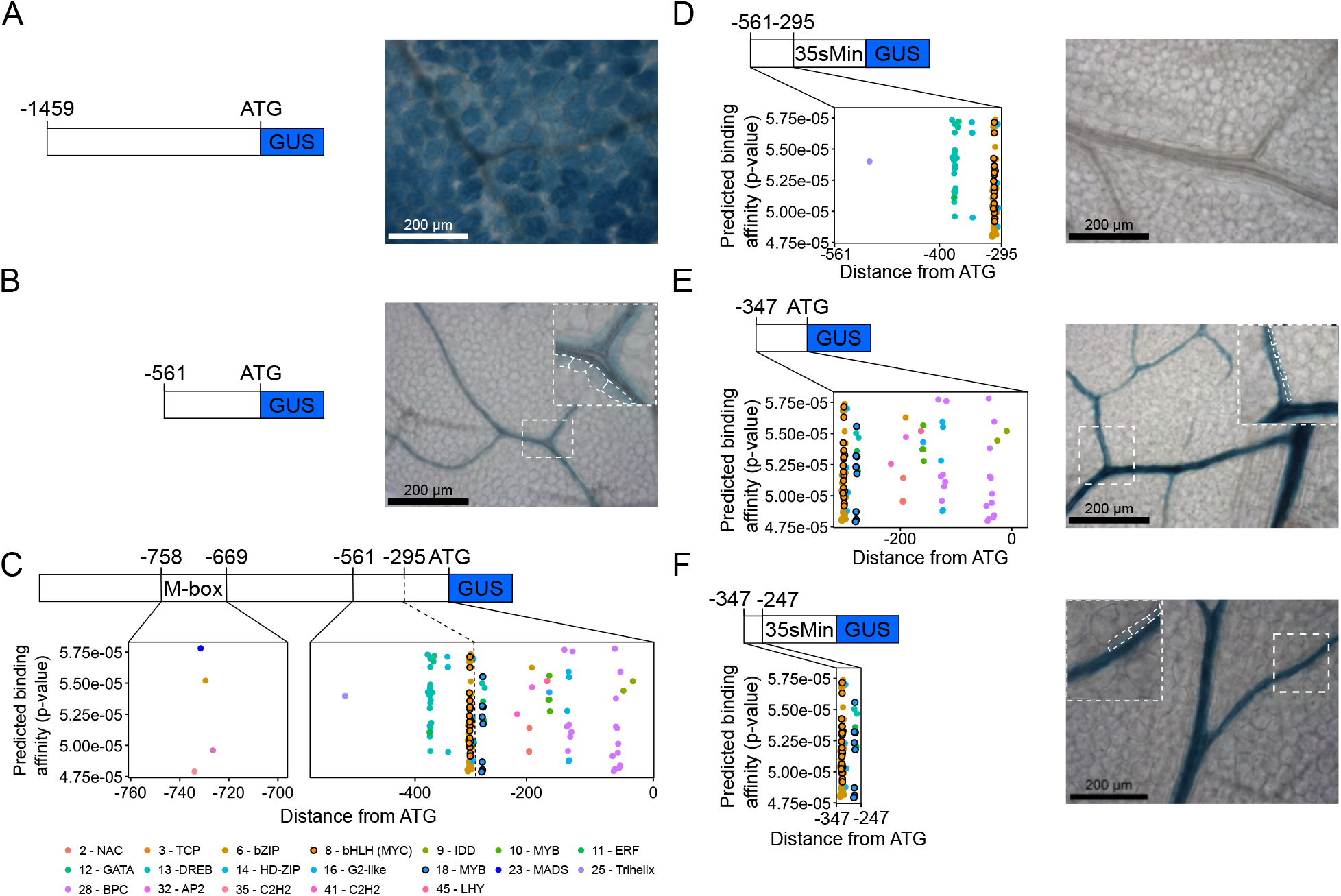
MYC and MYB TF binding motifs control bundle sheath strand expression in *Arabidopsis thaliana*. 1A) Schematic and representative GUS staining image of the full length -1458 bp *A. thaliana GLDP1* promoter upstream of the translational start site (ATG) from 19 independent T1 lines. B) Schematic and representative GUS staining image of the -561 bp *A. thaliana GLDP1* promoter upstream of the ATG from 13 independent T1 lines C) Predicted TF binding motifs in the M-box and from -561 bp upstream to the ATG of the *A. thaliana GLDP1* promoter. The position in the promoter (bp) is on the x axis, and the predicted binding affinity (p-values calculated from the log-likelihood score by the FIMO tool is on the y axis). The motifs are coloured by the motif clusters shown underneath the plots (Supplemental Table 1). D to F) Transcription factor binding motifs and representative GUS staining images of nucleotides - 561 to -295 bp upstream of the ATG fused to *CaMV35sMin* (D), -347 bp upstream to the ATG (E) and -100 bp sequence from -347 to -247 bp upstream of the translational start site fused to *CaMV35sMin* (F) from 17, 7 and 14 independent T1 lines respectively. Distance from the ATG (bp) is on the x-axis, and the predicted binding affinity (p-values calculated from the log-likelihood score by the FIMO tool (Grant et al., 2011)) is on the y axis. On GUS images, leaves were stained for 24 (A, B, E, and F) or 48 (D) hrs, scale bars are 200 μm and a zoom in of a region of the image is marked by a dashed white box. Bundle sheath cells marked with a dashed white box on the zoomed in inlay.

To better understand mechanisms controlling the expression of *GLDP1* in bundle sheath strands of *A. thaliana* we first used publicly available data to identify potential transcription factor binding sites in the promoter (Fig. 1C). Because not all transcription factor binding sites have been defined and some transcription factors are predicted to bind to the same or very similar sequences, we clustered motifs from the JASPAR database (Fornes, 2020) by similarity (Supplemental table 1). This provided an indication of transcription factor families likely able to bind the *GLDP1* promoter. In the 59 base pair M box, binding sites for C2H2, MADS, bZIP and BPC transcription factor families were present (Fig. 1C) suggesting that members of these families could be responsible for generating expression in the mesophyll. Although previous work had shown that nucleotides -561 to -295 upstream of the translational start site of *GLDP1* are necessary for expression in bundle sheath strands (Adwy et al., 2015) it is not known if they are sufficient for this patterning. We therefore searched for transcription factor binding sites in this region but also in the sequence up to the translational start site (Fig. 1C). Motifs associated with sixteen families of transcription factor families were identified, and this included closely spaced MYELOCYTOMATOSIS (MYC - belonging to the bHLH family) and MYOBLASTOMA (MYB) binding sites (Fig. 1C).

A bipartite module involving MYC2,3&4 and MYB28&29 directs expression of the *MYB76* transcription factor and glucosinolate biosynthesis genes to the bundle sheath of *A. thaliana* (Dickinson and Knerova et al., 2020). Re-analysis of publicly available data showed that *GLDP1 ex*pression was not reduced in leaves of the triple my*c2/3/4* mutant (Major et al., 2017). However, in the double *myb28/29* mutant (Burow et al., 2015) a small reduction in *GLDP1* transcript abundance was apparent (Supplemental Fig. 1B). As the bundle sheath of *A. thaliana* comprises only ∼15% of all cells in the leaf (Kinsman and Pyke, 1998), expression of *GLDP1* in the mesophyll will dominate signal from whole leaves. We thus considered this small change in *GLDP1* expression in the *myb28/29* mutant allele as consistent with MYB transcription factors controlling *GLDP1* expression in the bundle sheath. We hypothesised that the closely spaced MYC and MYB motifs drive expression of *AtGLDP1* in bundle sheath strands that becomes easily detectable once function of the mesophyll box is lost. Loss of the MYC binding site between nucleotides -305 and -299 upstream of the ATG abolished expression in bundle sheath strands (Adwy et al., 2015). However, the importance of of the MYB site between nucleotides -284 and -277 has not been investigated (Fig. 1C). We therefore conjectured that the region containing only the MYC binding site would not be sufficient for expression in bundle sheath strands. Consistent with this, when nucleotides -561 to -295 were fused to the minimal *CaMV35S* promoter, GUS activity was not detected (Fig. 1D, Supplemental Fig. 4). In contrast, when the MYC and MYB binding sites were both present, bundle sheath expression was restored (Fig. 1E, Supplemental Fig. 5). This indicates that sequence upstream of the MYC binding site is not necessary for expression in bundle sheath strands. When only the MYC and MYB binding sites were fused to the minimal *CaMV35S* promoter, GUS activity was detected in bundle sheath strands (Fig. 1F, Supplemental Fig. 6). We conclude that closely spaced MYC and MYB binding sites, from nucleotide -305 to -277, in the C_3_ *A. thaliana GLDP1* promoter are necessary and sufficient for bundle sheath strand expression. Thus, this MYC-MYB module can act alone to generate expression of genes such as *MYB76* in bundle sheath strands (Dickinson and Knerova et al., 2020) but as seen for *AtGLDP1*, it can also act in concert with other elements such as the mesophyll box to ensure expression in both bundle sheath and mesophyll cells. We next sought to test whether the MYC-MYB module is conserved in *GLDP1* genes from C_2_ species.

### C_2_ *Moricandia* species contain conserved and functional MYC and MYB binding sites in the *GLDP1* promoter

The Brassicaceae contains at least five independent origins of C_2_ photosynthesis (Schlüter et al., 2023). We hypothesized that conservation of MYC and MYB binding sites driving bundle sheath expression of *GLDP1* across the Brassicaceae underpins the repeated evolution of this trait. To test this, we aligned *GLDP1* promoter sequences from 17 species across the Brassicaceae including nine C_3_ and eight C_2_ species representing the five independent origins of C_2_ photosynthesis (Guerreiro et al., 2023). The MYC binding site (*CACGTG*) is perfectly conserved in all 17 species analysed and the MYB binding site (*CACCAAC*) is perfectly conserved in all species except *B. gravinae* and *D. tenuifolia* where a single substitution at position five of the motif replaced thymine with adenine (Fig. 2A). This suggests that the MYC and MYB binding sites responsible for driving bundle sheath strand expression of *GLDP1* in *A. thaliana* may be functional across these C_3_ and C_2_ Brassicaceae species. These data indicate that *cis*-elements allowing expression in bundle sheath strands have remained stable for at least 20.8 Ma since the divergence of *Arabidopsis* and *Moricandia* (Schlüter et al., 2017).

**Figure 2.**
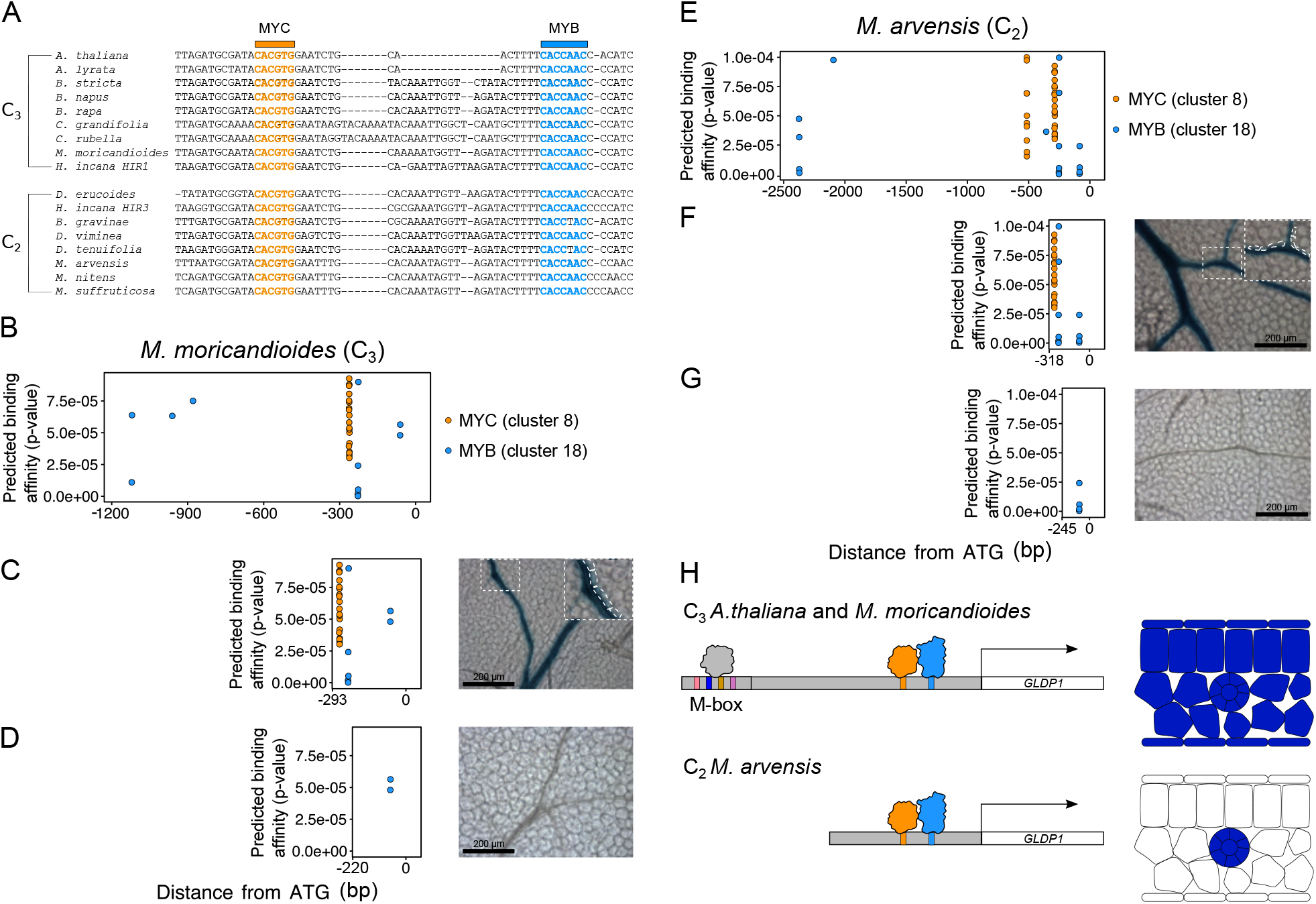
MYC and MYB binding sites are conserved in the Brassicaceae and drive vein and bundle sheath preferential expression of *Moricandia GLDP1* genes. A) Sequence alignments of the region of Brassicaceae *GLDP1* promoters containing MYC and MYB TF binding sites. MYC and MYB TF biding sites are coloured in gold and blue and marked above the alignment. B) Position of MYC and MYB binding sites in the *M. moricandioides GLDP1* promoter. C) Position of MYC and MYB binding sites and representative GUS staining images from 18 and 11 independent T1 lines respectively for *M. moricandioides* -293 bp and (D) -220 bp promoters. E) Position of MYC and MYB binding sites in the *M. arvensis GLDP1* promoter. F) Position of MYC and MYB binding sites and representative GUS staining images from 19 and 9 independent T1 lines respectively for *M. arvensis* -318 bp and (G) -245 bp promoters. Distance from the ATG (bp) is on the x axis, and the predicted binding affinity (P values calculated from the log-likelihood score by the FIMO tool (Grant et al., 2011) is on the y axis. On GUS images, leaves were stained for 24 (C and F) or 48 (D and G) hrs scale bars are 200 μm and a zoom in of a region of the image is marked by a dashed white box. Bundle sheath cells marked with a dashed white box on the zoomed in inlay. H) Schematic showing model for the control of *GLDP1* expression in C_3_ *A. thaliana* and *M. moricandioides* (top) and C_2_ *M. arvensis* (bottom). In C_3_ species constitutive expression is driven by unknown transcription factor(s) activating mesophyll expression from the M box (Adwy et al., 2015), potentially through binding to motifs from C2H2, MADS, bZIP and/or BPC families (Fig. 1C), and MYC and MYB TFs binding to closely spaced TF binding motifs to activate expression in the vein and bundle sheath. In C_2_ species the M-box is unable to activate expression in the mesophyll however MYC and MYB binding sites are conserved leading to bundle sheath strand specific expression of *GLDP1*.

To test whether these motifs are functional in Brassicaceae species in addition to *A. thaliana* we used the *Moricandia* genus for further investigation. *Moricandia* contains C_3_ and C_2_ species and previous work has shown that a promoter region, containing the predicted MYC and MYB binding sites is necessary for expression in the bundle sheath strand (Adwy et al., 2015; Adwy et al., 2019). To test whether MYC and MYB binding sites were able to drive expression in bundle sheath strands we cloned fragments from *GLDP1* promoters of C_3_ *M. moricandioides* and C_2_ *M. arvensis*. In C_3_ *M. moricandioides* closely spaced MYC and MYB sites are found between nucleotides -293 and -220 upstream of the predicted translational start site (Fig. 2B). Promoter deletions that removed all upstream sequence but retained the MYC and MYB sites, or also removed the MYC and MYB sites themselves were generated. When these motifs were present GUS activity was detected in bundle sheath strands (Fig. 2C, Supplemental Fig.7) but when they were absent this was not the case (Fig. 2D, Supplemental Fig. 8). Therefore, this C_3_ member of *Moricandia* contains sequence in the *GLDP1* promoter that is recognised by the MYC-MYB module of *A. thaliana* and it is able to pattern gene expression to bundle sheath strands. The *GLDP1* promoter from C_2_ *M. arvensis* also has closely spaced MYC and MYB motifs (Fig. 2E). When they were present, GUS activity was detected in *A. thaliana* bundle sheath stands (Fig. 2F, Supplemental Fig. 9) but when they are absent it was not (Fig. 2G, Supplemental Fig. 10).

Taken together, these data show that the bipartite MYC and MYB transcription factor module responsible for directing *MYB76* and glucosinolate biosynthesis genes to bundle sheath strands of *A. thaliana* is also used to pattern expression of *GLDP1* to this tissue. Moreover, the *cis*-code that is necessary and sufficient for bundle sheath strand expression is found in *GLDP1* genes from C_3_ and C_2_ species of *Moricandia*. The evolution of C_2_ photosynthesis in the Brassicaceae is thus associated with a shift of the M-box which disrupts its function (Triesch et al., 2022) and retention of closely spaced MYC and MYB binding sites such that *GLDP1* is expressed specifically in bundle sheath strands (Fig. 2H). Overall, this reveals a molecular genetic mechanism underpinning the bundle sheath accumulation of glycine decarboxylase required for C_2_ photosynthesis, and thus for a foundational step in the evolution of the C_4_ photosynthetic pathway. Further analysis will be required to establish whether other C_2_ and C_4_ lineages have made use of this MYC-MYB transcription factor module or whether evolution has convergently recruited other transcription factors to pattern genes to the bundle sheath.

## Materials and methods

### Plant materials and growth conditions

*A. thaliana* was grown on Levington F2 soil in growth chambers set at 20 °C, with a 16-hour-photoperiod with a light intensity of 150 μmol m^−2^s^−1^ photon flux density and 60% relative humidity.

### Transcription factor binding site prediction, sequence alignments, cloning and GUS assays

Motif clustering was performed on plant transcription factor motifs downloaded from JASPAR using the RSAT tool (Castro-Mondragon et al., 2017) as reported previously (Dickinson and Knerova et al., 2020). The FIMO tool (Grant et al., 2011) was used to scan DNA sequences for matches to *A. thaliana* transcription factor binding motifs found in the JASPAR motif database (Fornes, 2020). To account for input sequence composition, a background model was generated using the fasta-get-markov tool from the MEME suite (Bailey et al., 2009). FIMO was then run with the default parameters and a P value cut-off of 1 × 10^−4^.

Brassicaceae *GLDP1* promoter sequences were retrieved from phytozome (Goodstein et al., 2012) and promoters of *Moricandia* species were taken from Adwy et al., (2019). Sequences were aligned using MUSCLE (Edgar, 2004) with default settings and alignments visualised with the UGENE tool (Okonechnikov et al., 2012).

Promoter GUS constructs were assembled using the Golden Gate system (Weber et al., 2011). Arabidopsis promoter fragments were isolated from genomic DNA by PCR (primers in supplemental table 1) and cloned into level 0 modules (module information in supplementary table 2). *Moricandia* promoter fragments were initially amplified from genomic DNA using primers, adding a 5’ ClaI and a 3’ XbaI overhang. The amplified promoter sequences were subcloned into the pJET1.2 cloning vector using the Thermo Scientific CloneJET PCR Cloning Kit following the manufacturer’s instructions. *Moricandia* promoter fragments for Golden Gate cloning were then amplified from these pJET vectors and cloning into level 0 modules. Level 1 constructs were then assembled to fuse the promoter fragments with the *CaMV 35sMinimal* promoter were required, the GUS reporter and Nos terminator. Level 2 constructs were then assembled to add the FastR selectable marker (Shimada et al., 2010) to allow selection of positive transformants. Level 2 constructs were then placed into *Agrobacterium tumefaciens* strain GV3101 and introduced into *A. thaliana* Col-0 by floral dipping (Clough and Bent, 1998).

To take into account position effects associated with the transgene insertion site, GUS staining was undertaken on at least six randomly selected T1 plants for each *uidA* fusion. The staining solution contained 0.1 M Na_2_HPO_4_ (pH 7.0), 2 mM potassium ferricyanide, 2 mM potassium ferrocyanide, 10 mM EDTA (pH 8.0), 0.06% (v/v) Triton X-100 and 0.5 mg ml^−1^ X-gluc. Leaves from three-week-old plants were vacuum-infiltrated three times in GUS solution for one minute and then incubated at 37 °C for 24 h. Next, stained samples were fixed in 3:1 (v/v) ethanol:acetic acid for 30 minutes at room temperature, cleared in 70% (v/v) ethanol at 37 °C and then placed in 5 M NaOH for 2 h. The samples were stored in 70% (v/v) ethanol at 4 °C. The samples were imaged with an Olympus BX41 light microscope with Q Capture Pro 7 software and a QImaging MicroPublisher 3.3 RTV camera.

## Acknowledgements

The work was funded by the Advanced European Research Council Grant 694733 REVOLUTION, BBSRC grant BBW00013X1 to JMH and European Union Program (project GAIND4CROPS GA number 862087) to JMH and APMW. Work in the group of APMW was funded by ERA-CAPS project “C4BREED” under Project ID WE 2231/20–1, the Cluster of Excellence for Plant Sciences (CEPLAS) under Germany’s Excellence Strategy EXC-2048/1 under project ID 390686111, and the CRC TRR 341 “Plant Ecological Genetics” grant by the German Research Foundation (DFG). For the purpose of open access, the authors have applied a Creative Commons Attribution (CC BY) license to any Author Accepted Manuscript version arising from this submission.

