## Supplemental Figures for "A transcription factor module mediating C_2_ photosynthesis"

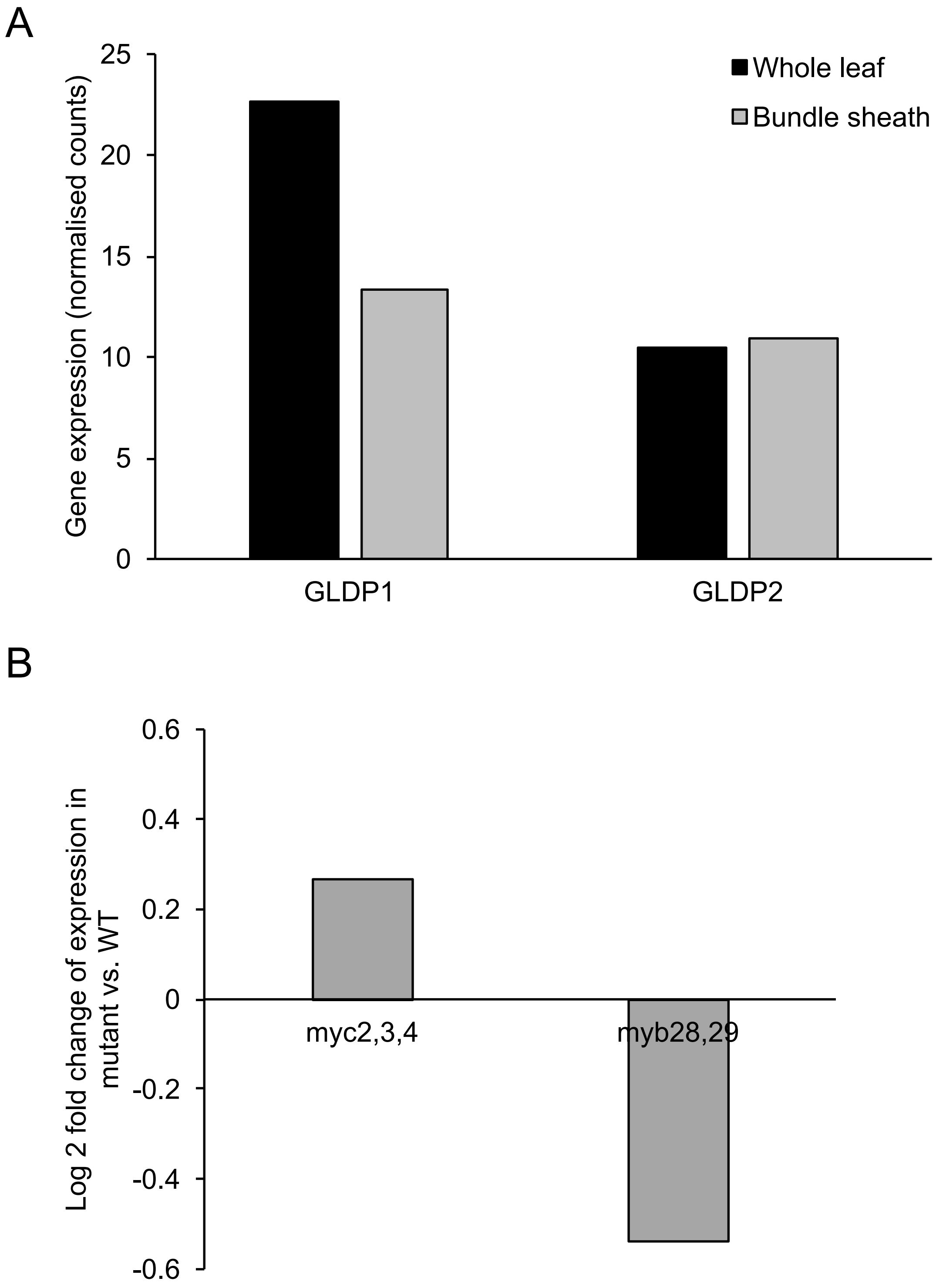


**Supplemental Figure 1. Expression of *A. thaliana GLDP* genes. A)** Expression of *AtGLDP1* and *AtGLDP2* in whole leaf and bundle sheath translatomes (Aubry et al., 2013). **B)** Log2 fold change of *AtGLDP1* expression in *myc,2,3,4* mutants (Major et al., 2017) and *myb28.29* mutants (Burow et al., 2015) vs. WT.

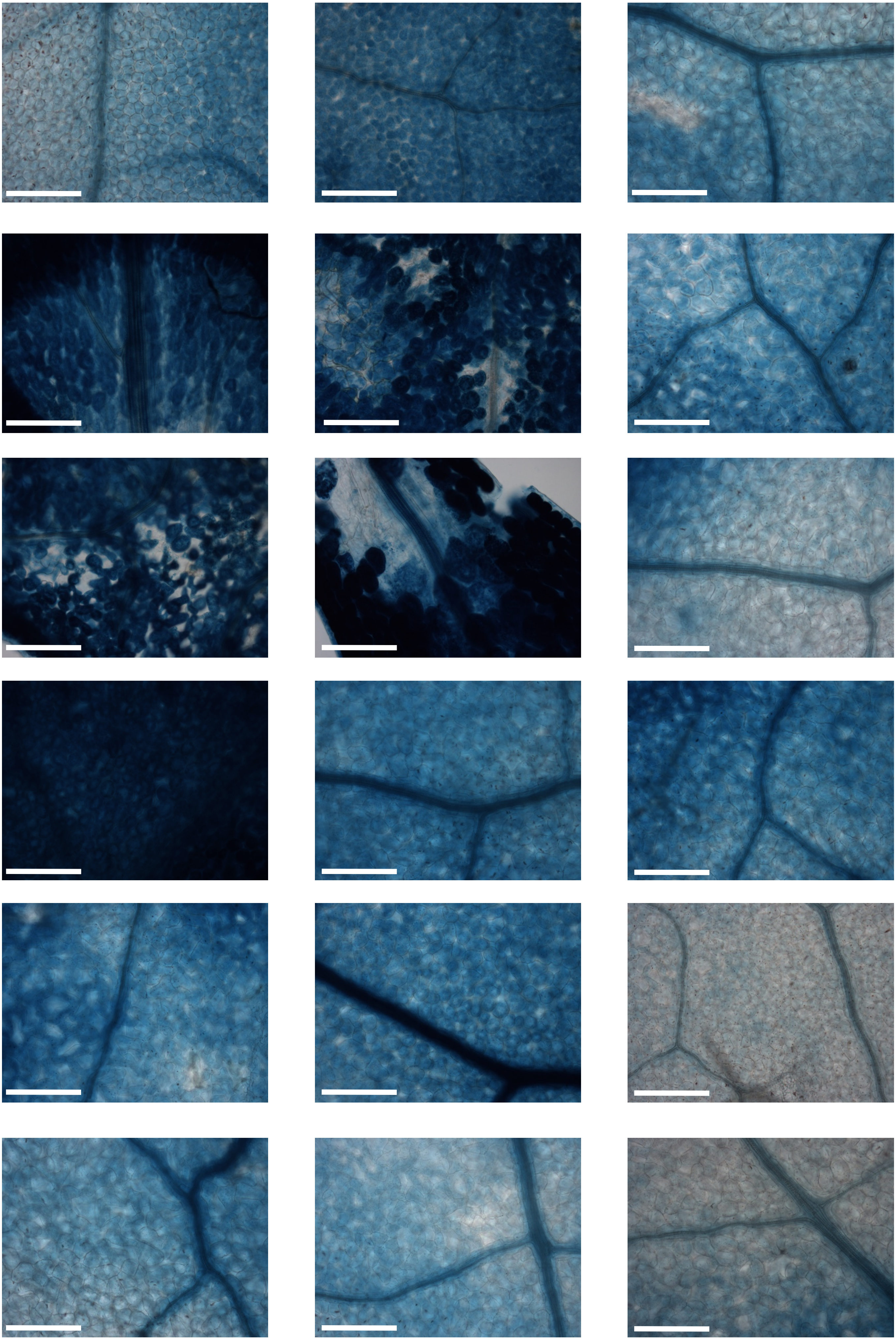

**Supplemental Figure 2.** Nucleotides -1458 bp to the translational start site (ATG) of *AtGLDP1* generate expression in the bundle sheath and mesophyll. Images from 18 independent transgenic lines. Leaves were stained for 24 hrs. Scale bars represent 200 µm.

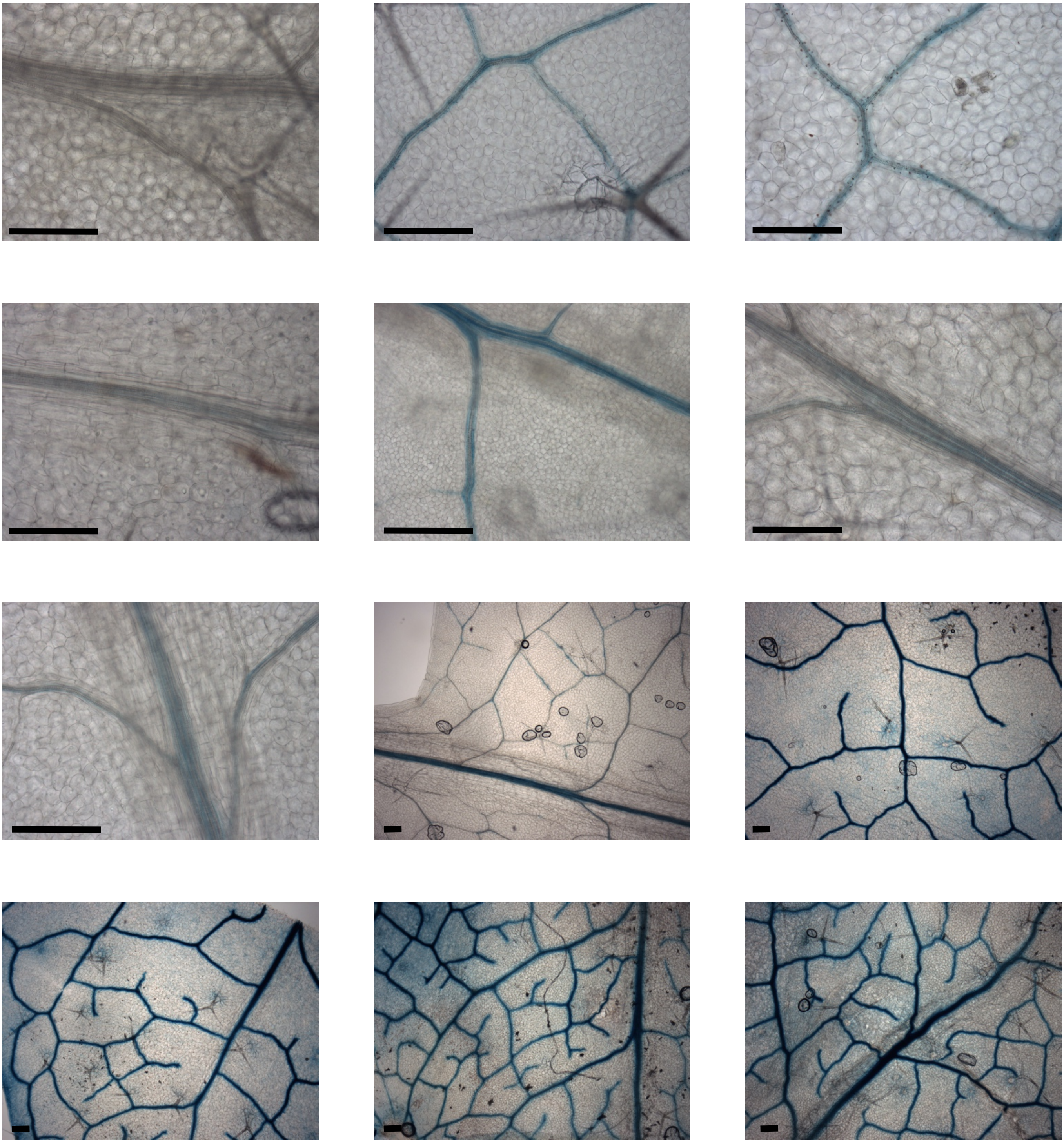

**Supplemental Figure 3.** Nucleotides -561 bp to the ATG of *AtGLDP1* drive expression in the bundle sheath strand. Images from 12 independent transgenic lines. Leaves were stained for 24 hrs. Longer scale bars represent 200 µm, shorter scale bars represent 20 µm.

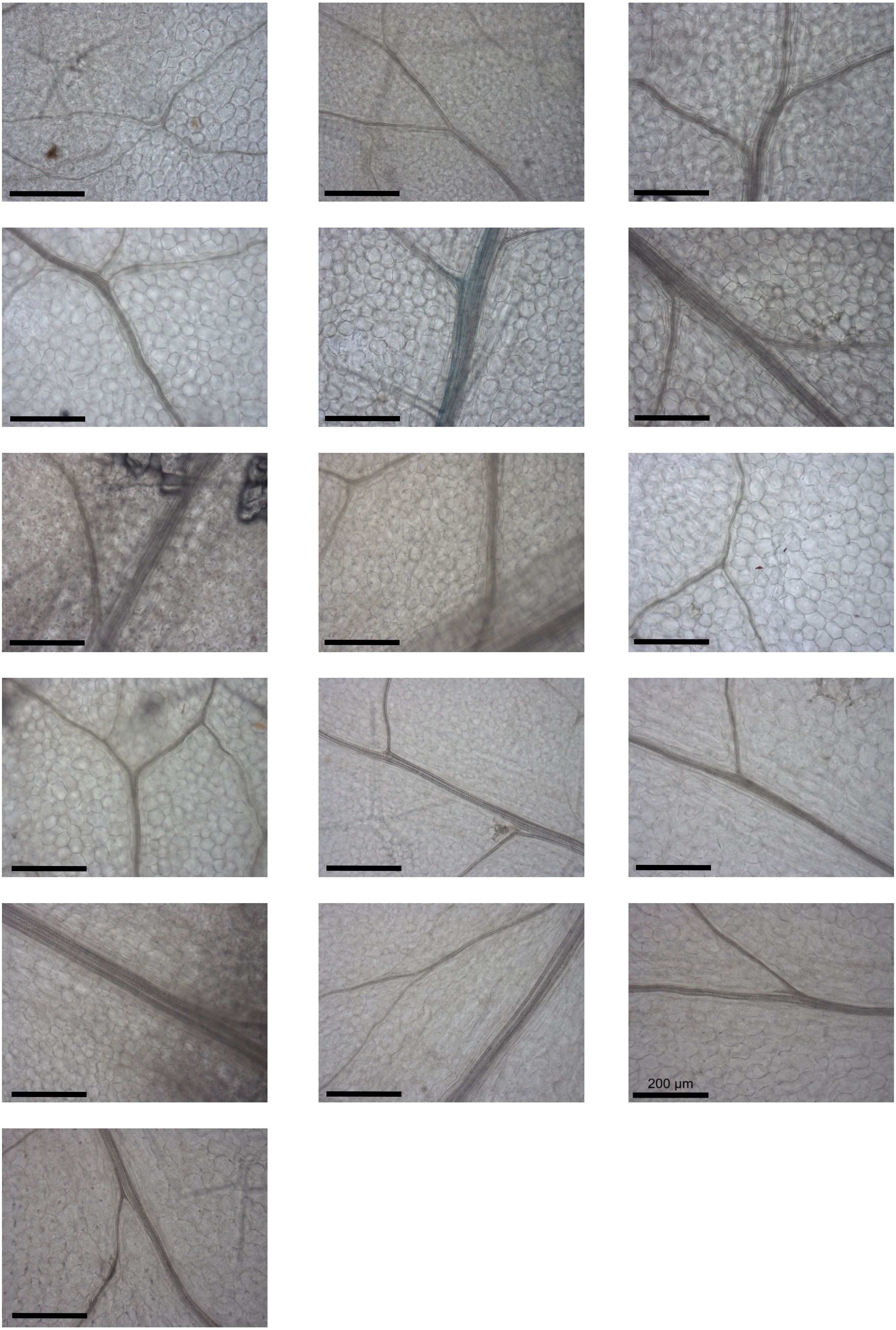
200 µm

**Supplemental Figure 4.** Nucleotides -561 to -295 bp upstream of the ATG of *AtGLDP1* fused to CaMV35sMin do not drive expression in the bundle sheath strand. Images from 16 independent transgenic lines. Leaves were stained for 48 hrs. Scale bars represent 200 µm.

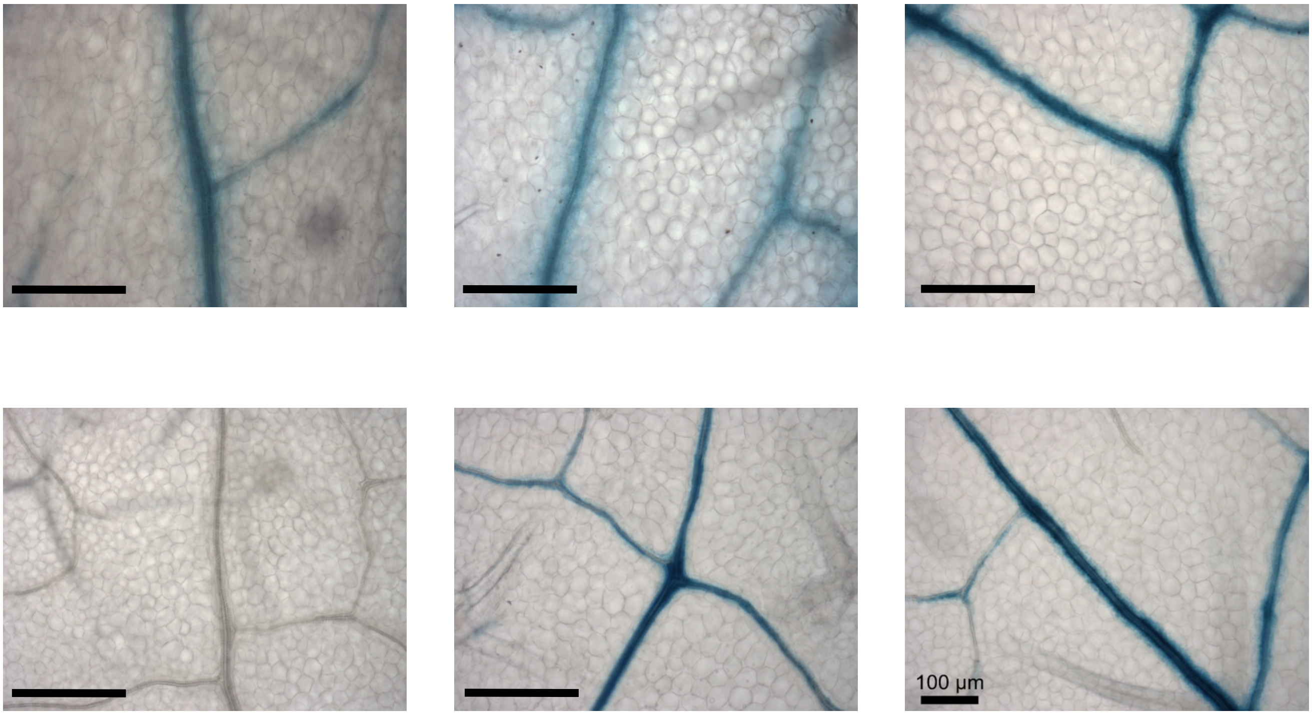


**Supplemental Figure 5.** Nucleotides -347 bp upstream to the ATG of *AtGLDP1* can drive expression in the bundle sheath strand. Images from 6 independent transgenic lines. Leaves were stained for 24 hrs. Scale bars represent 200 µm.

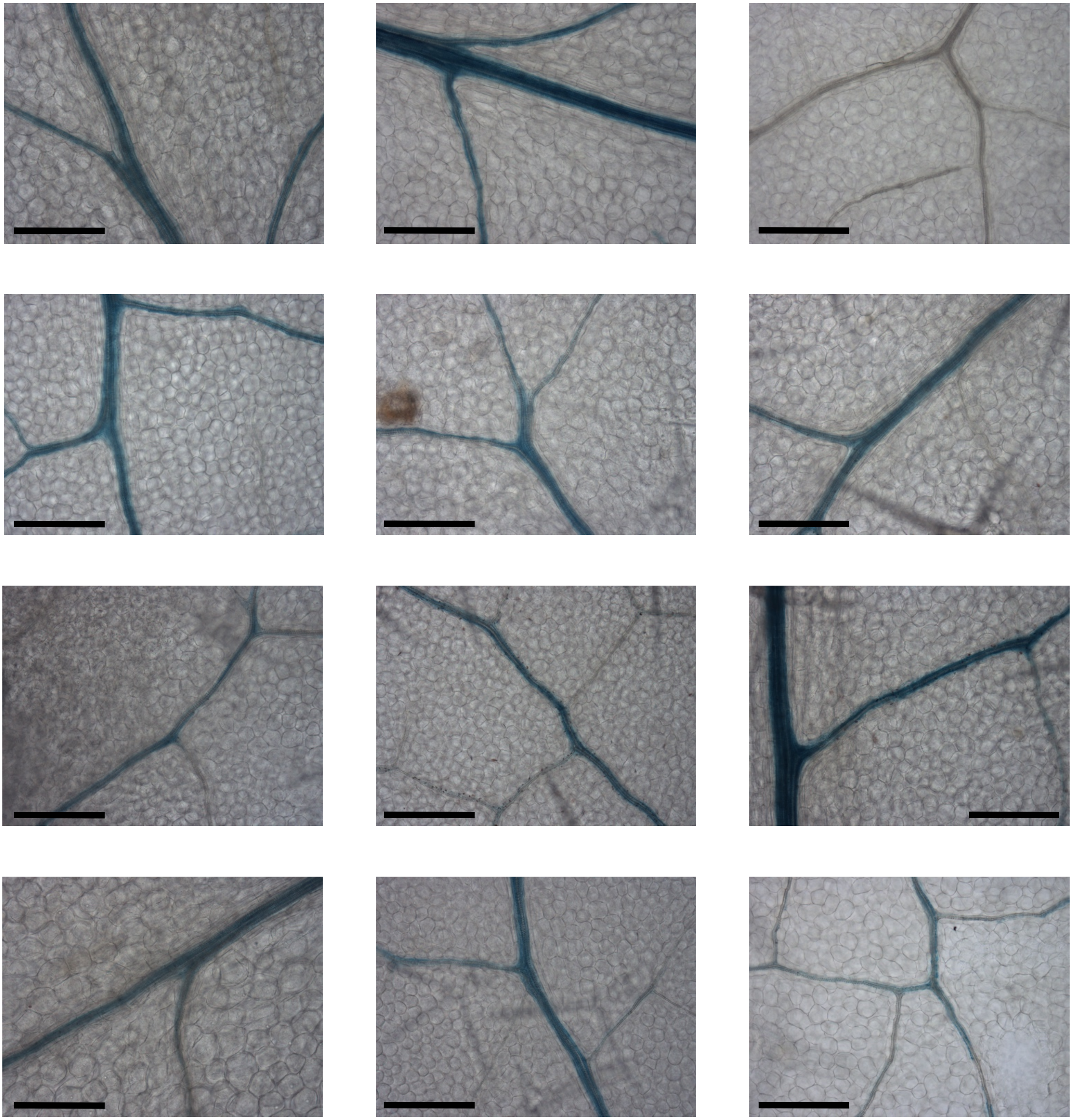

**Supplemental Figure 6.** 100 bp sequence from -347 to -247 bp upstream of the ATG of *AtGLDP1* fused to CaMV35sMin can drive expression in the bundle sheath strand. Images from 12 independent transgenic lines. Leaves were stained for 24 hrs. Scale bars represent 200 µm.

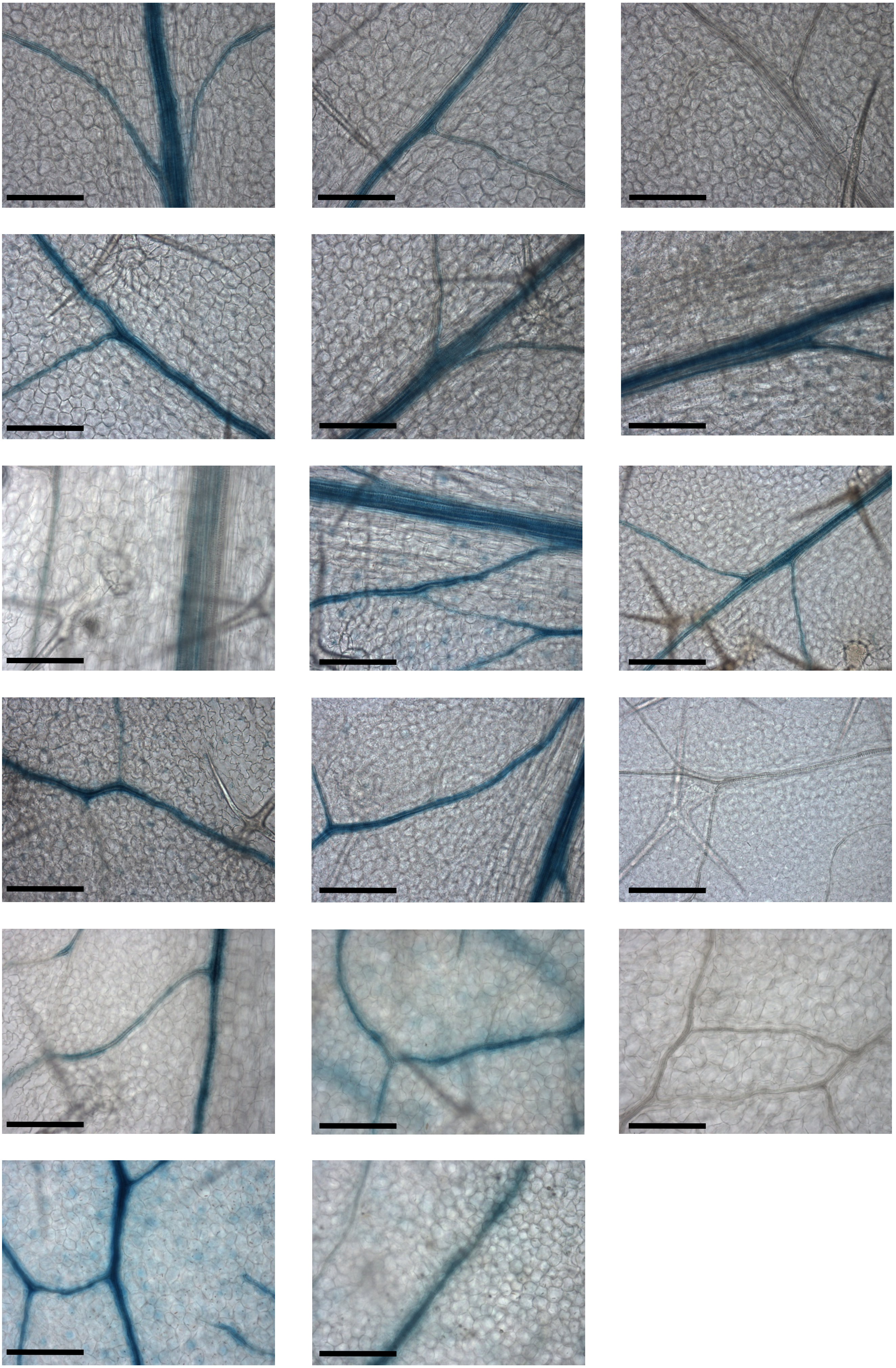

**Supplemental Figure 7.** Nucleotides from -293 bp upstream to the ATG of *M. moricandioides GLDP1* can drive expression in the bundle sheath strand. Images from 17 independent transgenic lines. Leaves were stained for 24 hrs. Scale bars represent 200 µm.

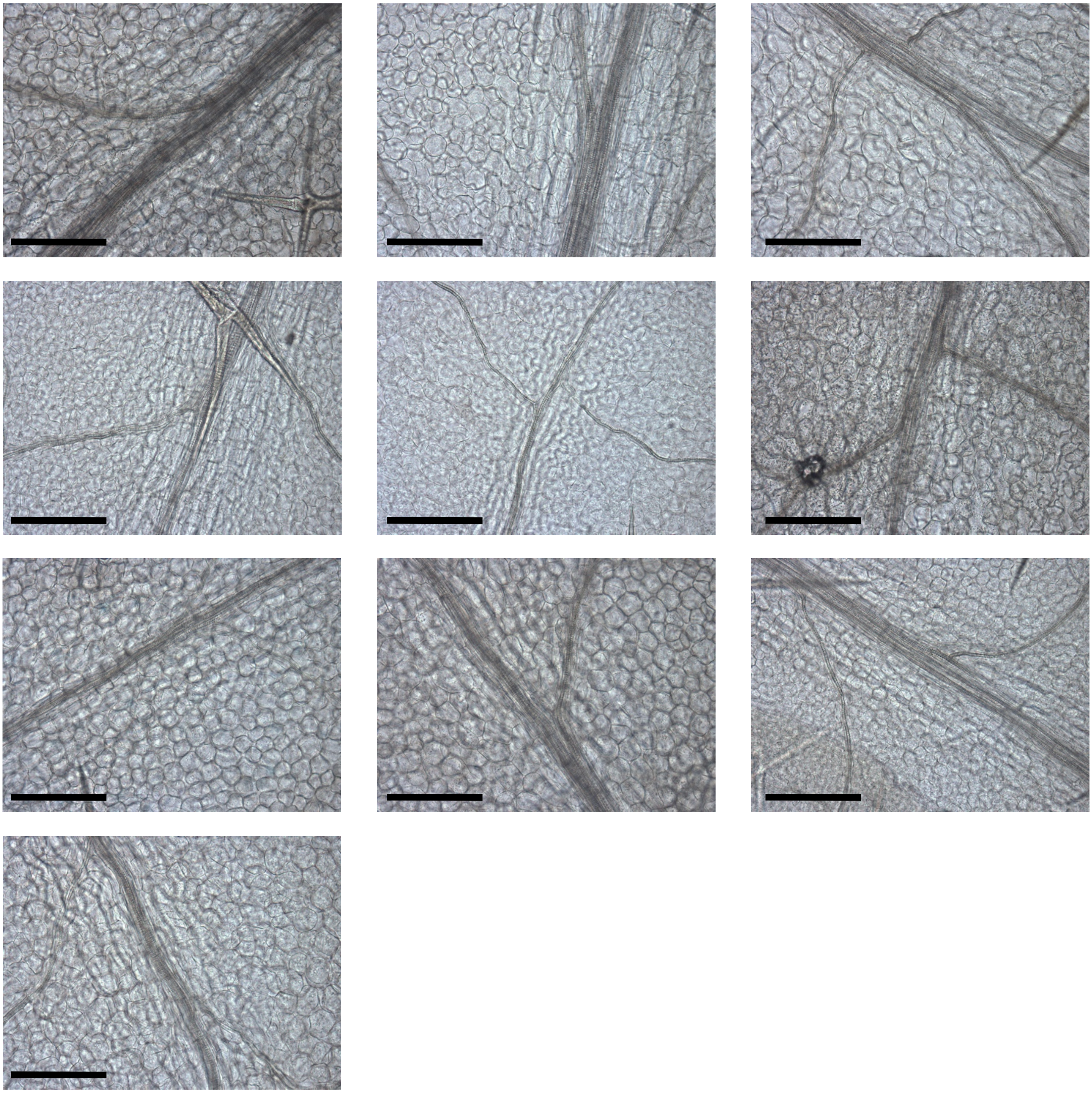

**Supplemental Figure 8.** Nucleotides from -220 bp upstream to the ATG of *M. moricandioides GLDP1* do not drive expression in the bundle sheath strand. Images from 10 independent transgenic lines. Leaves were stained for 48 hrs. Scale bars represent 200 µm.

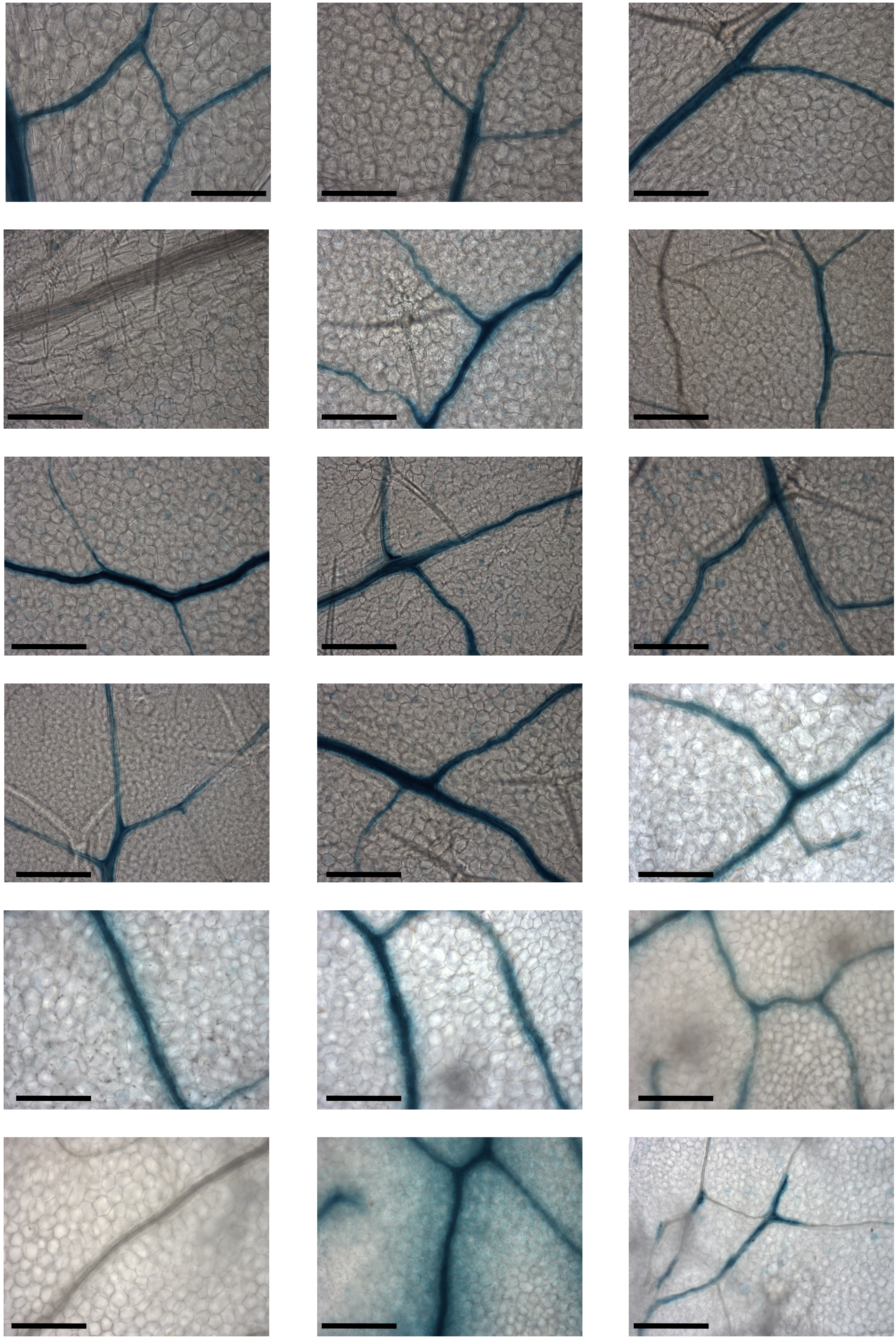

**Supplemental Figure 9.** Nucleotides from -318 bp upstream to the ATG of *M. arvensis GLDP1* can drive expression in the bundle sheath strand. Images from 18 independent transgenic lines. Leaves were stained for 24 hrs. Scale bars represent 200 µm.

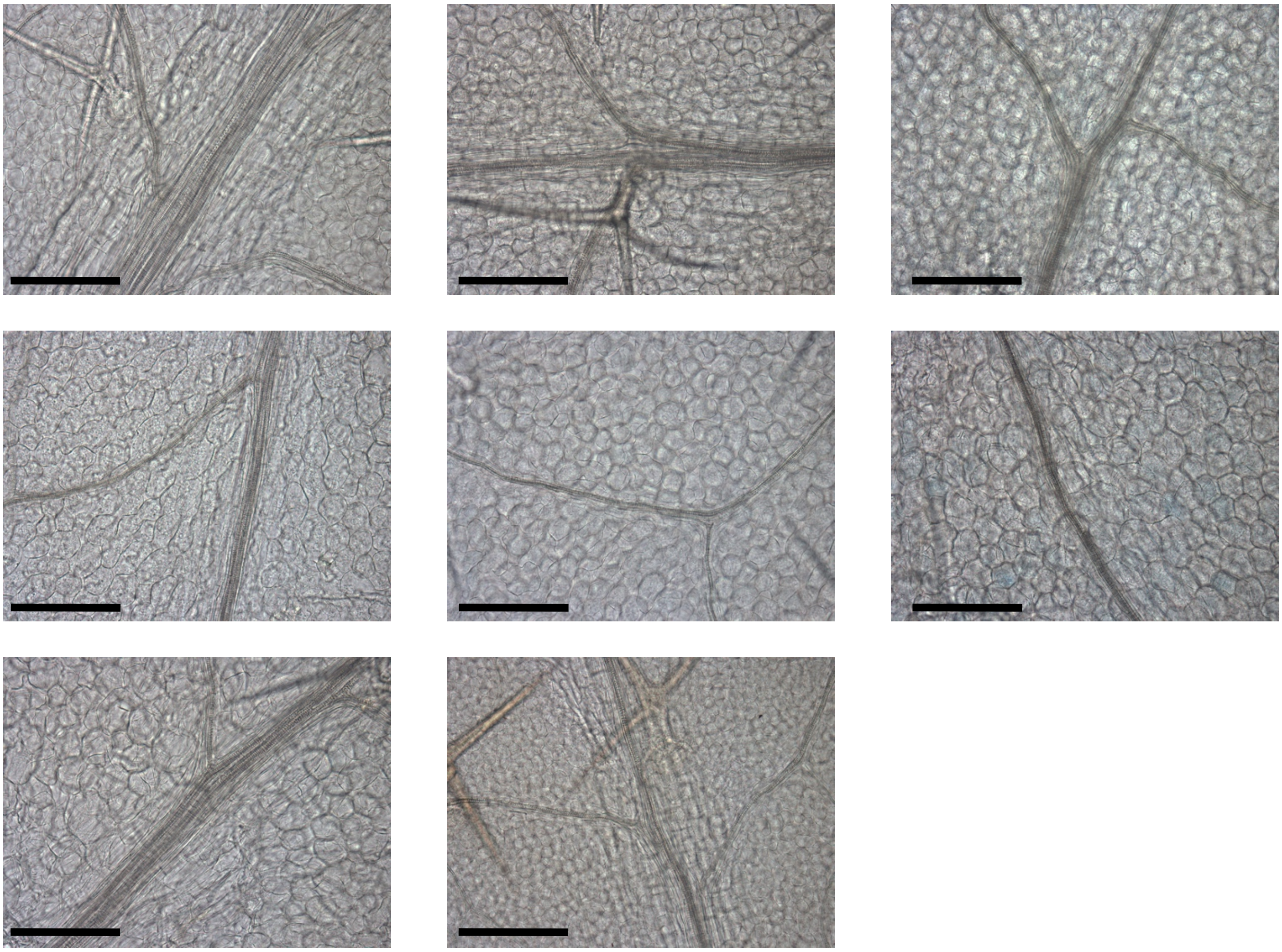

**Supplemental Figure 10.** Nucleotides from -245 bp upstream to the ATG of *M. arvensis GLDP1* do not drive expression in the bundle sheath strand. Images from 10 independent transgenic lines. Leaves were stained for 48 hrs. Scale bars represent 200 µm.
